# Emergence of meniscus-guided movement in drosophilid larvae through posture-dependent capillary forces

**DOI:** 10.64898/2026.05.03.722455

**Authors:** Teruyuki Matsunaga, Akinao Nose

## Abstract

Freshwater habitats cover only ∼1% of the Earth’s surface yet harbor approximately 10% of all animal species, of which ∼60% are insects, making them hotspots of biodiversity. However, tractable model systems to investigate how insects transition to aquatic environments remain limited. Here, we show that larvae of *Scaptodrosophila dorsocentralis*, but not related species including *Drosophila melanogaster*, move along the water meniscus by exploiting surface tension, enabling them to reach nearby objects. This movement is achieved through a sequence of actions: larvae adopt an S-shaped body posture by extending the posterior body, anchor at the air-water interface, and generate propulsive forces by elevating the anterior end while depressing the posterior end. Larvae successfully reach and land on nearby objects via meniscus-guided movement even under flowing conditions, whereas other species fail to do so, indicating ecological relevance. A biomimetic PDMS (polydimethylsiloxane) model recapitulates this movement without external actuation, demonstrating that body configuration alone is sufficient to generate capillary-driven motion. We further show that posterior elongation is mediated by a folding–unfolding mechanism driven by hydrostatic pressure. These results establish a tractable system for studying water-surface locomotion and provide mechanistic insight into how terrestrial insects may acquire the capacity to exploit water-surface environments.

## Main

Aquatic habitats cover only ∼1% of the Earth’s surface yet harbor approximately 10% of all animal species, of which ∼60% are insects ^1,2^. Aquatic insects play key ecological roles in freshwater ecosystems, including nutrient cycling and serving as a major food source for higher trophic levels such as fish ^3^. Thus, their diversity represents not only a central subject in studies of biodiversity and speciation but also a critical component of freshwater ecosystem health. Identifying the mechanistic bases of transitions to aquatic habitats is therefore essential for understanding how insects evolve across ecological niches, acquire novel ecological functions, and contribute to the maintenance of healthy environments^4,5^.

For small terrestrial insects, however, water presents substantial physical challenges^6^. At the millimetre scale, viscous forces and surface tension dominate over inertia, making movement at the air–water interface difficult. For example, even a raindrop can trap insects, posing a serious threat to survival and fitness ^7,8^. While many aquatic or semi-aquatic insects, such as water striders, firefly larvae, and mayflies, have evolved specialized morphologies and behaviors for locomotion on or within water, the mechanisms by which terrestrial insects initially adapt to these environments remain poorly understood ^9,10^. Two major limitations have hindered progress in this area. First, many aquatic insect lineages diverged hundreds of millions of years ago, and extensive neutral sequence divergence obscures causal genotype–phenotype relationships^11^. Second, most aquatic insects are not easily maintained under laboratory conditions, limiting functional and mechanistic analyses.

Previous studies have shown that a range of insects, including semi-aquatic species such as water treaders and water lily leaf beetles, can exploit the upward meniscus formed at the interface between water and hydrophilic objects to climb surfaces ^12–14^. This behavior represents a potential intermediate step toward aquatic adaptation.

However, such species are not readily amenable to genetic manipulation, and comparable behaviors have not been reported in tractable laboratory systems such as drosophilids. Establishing a system that combines ecological relevance with experimental accessibility is therefore critical for dissecting the mechanisms underlying transitions to aquatic environments.

Diptera contains the largest number of aquatic species among insects, with approximately one-third (∼46,000 species) utilizing aquatic habitats during at least one life stage ^1,3^. Many of these transitions appear to have evolved independently, suggesting that shifts from terrestrial to aquatic environments may be relatively recent and evolutionarily accessible^15^. Importantly, Diptera also includes genetically tractable model organisms such as *Drosophila melanogaster*, which has been studied for over a century^16^. This provides an opportunity to identify related systems in which early stages of aquatic adaptation can be investigated by leveraging existing genetic knowledge and partially transferable experimental frameworks. However, to our knowledge, meniscus-climbing behavior, a potential early step in semi-aquatic adaptation, has not been reported in Diptera. This gap has limited our ability to understand how the initial stages of aquatic adaptation evolve in insects.

Here, we report that larvae of *Scaptodrosophila dorsocentralis*^17^, a lineage restricted to the islands of Okinawa, Japan, exhibit meniscus-guided movement, whereas closely related species including *S. coracina, Drosophila melanogaster,* and *Chymomyza procnemis* do not^18–20^. This enables larvae to move along the water surface and reach nearby objects, even under flowing conditions. Movement is achieved through a coordinated sequence of behavioral components, including tail elongation, maintenance of an S-shaped body posture, and the exploitation of capillary forces. Quantitative analysis, based on established physical models ^21^, revealed that the magnitude of these forces is comparable to those reported in semi-aquatic insects, and this movement can be recapitulated using a PDMS-based physical mimic. Furthermore, tail elongation in *Scaptodrosophila* is mediated by a previously undescribed folding–unfolding mechanism driven by compression of the anterior-to-middle body segments, generating hydrostatic pressure. Together, our study establishes a tractable model system for investigating water-surface movement and provides mechanistic insights into how insects may acquire the capacity to exploit water-surface environments.

## MATERIAL AND METHODS

### Fly Husbandry

*Scaptodrosophila dorsocentralis* (IR-J02), *S. bryani* (KMJM25), *S. coracina* (IR-M3), *Chymomyza procnemis* (2631.01-TC), and *Drosophila mojavensis* (CI 12 IB-4 g8) were obtained from Kyorin-Fly. *Drosophila melanogaster* (Canton-S) was obtained from the Kyoto Stock Center, and *S. latifasciaeformis* was obtained from National Drosophila Species Stock Center (Cornell University). All species were reared on standard banana-based medium (recipe available on the Kyorin-Fly website) under a 12 h light/12 h dark cycle at >60% relative humidity. *D. melanogaster, D. mojavensis*, *S. dorsocentralis, S. bryani, S. latifasciaeformis*, and *C. procnemis* were maintained at 23 °C, whereas *S. coracina* was maintained at 18 °C due to reduced reproductive performance at higher temperatures.

### Larval landing assay on static water

A 7 cm-long, 2 cm-diameter branch of *Zelkova serrata* collected on campus was half-submerged in tap water in a 5 cm cubic glass chamber filled to a depth of 3 cm, generating an upward meniscus at the interface between the water surface and the branch. The setup is described in Supplementary Fig. 1A. Third instar larvae were gently rinsed with distilled water and dried using Kimwipes. Individual larvae were placed on the water surface with a paint brush at a distance of ∼1.5 cm from the branch. Larval trajectory on the water surface was recorded for 5 min from the dorsal view using a stereomicroscope (Olympus SZX16) equipped with a camera (Hozan L-836). For lateral recordings, a digital camera (Olympus Tg-6) in macro mode was used. Water was replaced between species to minimize potential chemical carryover effect. For anesthesia treatment, *S. dorsocentralis* larvae were exposed to CO2 for 1 min. Landing success was defined as reaching and contacting the branch via movement on the water surface within 5 min. Larvae that sank within 30 s of introduction were excluded from analysis.

### Larval landing assay on flowing water

A peristaltic pump (MINIPULS 3, Gilson, France) was used to generate water flow. Detailed dimensions of the setup are provided in Supplementary Fig. 1B. The arena was constructed from polyethylene, and the bottom near the water inlet and all side walls were lined with thin wooden tape to increase hydrophilicity and promote laminar flow. Movements of third instar larvae on the water surface were recorded from above using a camera (Mightex CGE-B013-U) at 1 fps for 10 min or until larvae reached the arena walls. Flow rates were controlled by setting the pump rotation speed to 12, 24, 36, or 48 rpm. The maximum flow rate (48 rpm) corresponded to approximately 29 mL min⁻¹, according to the manufacturer’s specifications. Before each assay, the arena was filled with approximately 250 mL of tap water, and the walls were gently wetted using a paintbrush to generate an upward meniscus. Third instar larvae were rinsed with distilled water and dried using Kimwipes, then gently placed on the water surface. Landing success was defined as reaching and contacting the arena wall via movement on the water surface within 10 min. Larvae that sank within 30 s of introduction were excluded from analysis. Total displacement along the x-axis (mm) and reaching time (s) were calculated only for individuals that successfully reached the arena wall; individuals that did not reach the wall were excluded from these analyses.

### Calculation of transition probability and S-shape posture duration

Third instar larvae were rinsed with distilled water, gently dried using Kimwipes, and individually placed on the surface of water in a glass cube. Larval behavior at the air–water interface was recorded laterally using a digital camera (Olympus TG-6) in macro mode.

Behavioral states were defined as follows.

### Surface

The posterior tip and at least one additional body region are in contact with the water surface.

### Dip

The anterior–middle part of the body detaches from the surface and moves downward into the water.

### Elevate

The direction of head movement reverses from downward to upward, and the anterior tip moves toward the surface.

### S-shape

The anterior tip, middle region, and posterior end are simultaneously in contact with the surface, and the body forms a laterally curved S-shaped configuration, with the posterior end positioned between the anterior and middle regions.

For quantification of S-shaped posture duration, larval behavior was recorded from a dorsal view using a camera (Mightex CGE-B013-U). The onset of the S-shaped posture was defined as the first frame in which the posterior tip was positioned in close proximity to the anterior body segments (approximately A2–A3), consistent with the S-shaped configuration. The end of the posture was defined as the frame in which this configuration was lost.

### Estimation of capillary forces from larval movement

The behavioral setup is described in Supplementary Fig. 1c. A total of 1.5 mL of distilled water was placed in a 3.5 cm-diameter culture dish (Thermo Scientific). Third instar larvae were rinsed with distilled water, dried using Kimwipes, and gently placed on the water surface. Movements were recorded from above using a camera (Mightex CGE-B013-U) at 37 fps for 2 min under infrared illumination using an IR-pass filter and two IR light sources (LDQ-150IR2-850, CCS).

Larval centroid displacements were extracted using LabGym^22^ after binarization in ImageJ, and velocities were calculated from these trajectories. Velocity as a function of time was fitted with an exponential function using nonlinear least-squares optimization in Python (scipy.optimize.curve_fit). Acceleration was obtained by analytical differentiation of the fitted function. The relationship between acceleration and displacement was then fitted with an exponential function of the form 𝑎(𝑥) = 𝑑𝑒 ^𝑥/𝑙𝑐^ + 𝑔, where 𝑙_𝑐_ = 0.27, using nonlinear least-squares optimization.

The tangential force balance was described as:

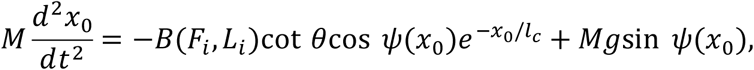

where

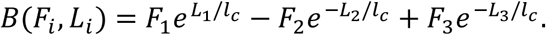

Acceleration–displacement relationships were fitted to estimate the parameter 𝐵, representing the combined capillary force contribution.

Force components 𝐹_1_, 𝐹_2_, and 𝐹_3_were then obtained by solving the following system of equations (normal force and torque balance):

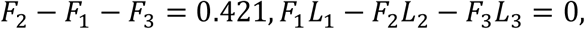

The system was solved numerically for each 𝐵using scipy.optimize.fsolve. Parameters were set to 𝜃 = 67.5^∘^, 𝑀𝑔 = 0.42, 𝐿_1_ = 0.1325, 𝐿_2_ = 0.007, 𝐿_3_ = 0.08585, and 𝑙_𝑐_ = 0.27. Values of 𝜃, 𝐿_1_, 𝐿_2_, and 𝐿_3_were measured from images using ImageJ. Body mass 𝑀was estimated as the mean mass of 200 flies measured using a precision balance (GX-200, A and D). Because the validity of the lateral force 𝐹, and thus the tangential force balance equation, is expected to break down when the meniscus slope becomes large, analyses were restricted to data points with 𝑥_0_greater than approximately 𝑙_𝑐_ = 0.27.

The constant term 0.421 was derived from 𝑀𝑔cos 𝜓(𝑥_0_)and represents an approximate value under the assumption 𝜓 ≈ 0in the region where larvae were sufficiently far from the wall.

### PDMS-based biomimetic assay

Polydimethylsiloxane (PDMS; Sylgard 184, Dow) was prepared according to the manufacturer’s instructions. The cured material was cut manually to approximate the S-shaped body of *S. dorsocentralis* larvae. The anterior and posterior ends of the PDMS mimic were connected using an insect pin (0.1 mm diameter, 12 mm length; 26002-10, Muromachi) to impose an asymmetric configuration, in which the anterior end was elevated and the posterior end was depressed, thereby generating upward and downward menisci, respectively. Control mimics were connected horizontally and did not produce apparent meniscus deformation (Fig. 4a).

A total of 1.5 mL of distilled water was placed in a 3.5 cm-diameter culture dish (Thermo Scientific). PDMS mimics and controls were individually placed on the water surface using forceps (INOX 5, Dumont). Movements were recorded from above using a camera (Mightex CGE-B013-U) at 37 fps for 2 min.

### Comparison of tail lengths across species

Agar (0.5%) was prepared for the dig-and-dive assay as previously described ^23^. Third instar larvae of each species were placed on the agar surface and allowed to burrow. During burrowing, tail length in contracted and extended states was recorded at 30 fps for 1 min using a stereomicroscope (Olympus SZX16) equipped with a camera (Hozan L-836), and measurements were performed using ImageJ. The tail-to-body length ratio was defined as the length from the posterior tip to the anus divided by the total body length (anterior tip to posterior tip). For each individual, the maximum tail length observed during the recording period was used for analysis.

### Visualization of folded tail structures in third instar *Scaptodrosophila* larvae

Third instar *S. coracina* larvae were rinsed with distilled water and dried using Kimwipes. The entire body was then coated with a commercially available green fluorescent dye using a paintbrush while the posterior end was in a contracted state. After 5–10 s, the tail was manually extended using forceps. Images before and after treatment were acquired using a stereomicroscope (Olympus SZX16) equipped with a camera (Hozan L-836). The non-fluorescent region length was defined as the length of the unlabeled posterior segment, and the non-fluorescent-to-body length ratio was calculated by dividing this length by the total body length (anterior tip to posterior tip).

### Visualization of folded tail structures in hatching embryos

Embryos of *D. melanogaster* were collected within a 0.5 h window at 19 h after egg laying (AEL) and fixed in 4% formaldehyde. To match the comparable developmental stages in *S. dorsocentralis*, embryos were collected within a 0.5 h window at 25 h AEL and fixed under the same conditions. Images were acquired from the lateral side using a stereomicroscope (Olympus, BX63) equipped with a 20× objective (Olympus, XLUMPlanFL N) under transmitted light at 647 nm. The tail-to-body area ratio was defined as the area of the folded posterior region divided by the total body area, measured using ImageJ.

### Tail compression and dynamics assay

Second instar larvae were fed banana-based medium supplemented with carbon black for 24 h prior to experiments to visualize the gut. On the following day, 0.5% agar was prepared for the dig-and-dive assay as previously described^23^. Third instar larvae with darkened intestines were placed on the agar surface and allowed to burrow. Images were acquired using a stereomicroscope (Olympus SZX16) equipped with a camera (Hozan L-836) at 30 fps for 1 min. Tail extension events that occurred within 1 s of the initiation of forward or backward peristaltic waves were excluded from analysis, as these movements can confound measurements of body length dynamics. Posterior length was defined as the distance between the posterior tip and the A7/A8 segment boundary, and anterior length as the distance between the anterior tip and the A1/A2 segment boundary (Supplementary Fig. 3). For each individual, the maximum change in posterior length within the recording period was used for analysis. Lengths were normalized within each individual using the maximum and minimum values observed in that animal. Time zero was defined as the point at which rapid tail shortening ceased. Cross-correlation analysis was performed between posterior and anterior length dynamics, with positive lag defined as posterior length changes preceding anterior length changes. Cross-correlation was computed using normalized signals.

## RESULTS

### Larval *Scaptodrosophila dorsocentralis*, but not closely related species, moves along the water meniscus and reaches nearby objects

We asked whether any drosophilid species can move at the air–water interface, a behavior not previously reported in this lineage, with the exception of a few fully aquatic larval species^24^. To mimic natural conditions, we placed an approximately 2 cm diameter and 7 cm length tree branch in a water-filled chamber (Supplementary Fig. 1A) for animals to land and tested multiple species across the genera *Scaptodrosophila, Drosophila*, and *Chymomyza*.

We found that larvae of *S. dorsocentralis*, but not other tested species including *S. bryani, S. latifasciaeformis, S. coracina, D. melanogaster, D. mojavensis,* or *C. procnemis*, exhibit a behavior resembling meniscus climbing previously described in semi-aquatic insects such as water lily leaf beetles, and water treaders^13,14,21^. The phylogenetic relationships among the tested species are shown in Fig. 1a. Specifically, *S. dorsocentralis* larvae anchor to the water surface via their hydrophobic posterior end^25^, adopt an S-shaped body posture, and move along the upward water meniscus to reach nearby objects (Fig. 1b, Supplementary Movie 1 and 2). Notably, larvae not only traverse the water surface but also reach and contact nearby objects and exit the water, indicating that this behavior enables effective escape from the air–water interface. In contrast, larvae of the other species also attach to the interface via the posterior end but exhibit uncoordinated head and tail movements without directional movement (Fig. 1c, Supplementary Movie 3). This meniscus-climbing behavior in *S. dorsocentralis* was abolished by CO₂ anesthesia, which disrupted formation of the S-shaped posture, indicating that this body configuration is essential for this movement (Fig. 1d; n = 10–38 animals per condition; Fisher’s exact test with Holm–Bonferroni correction, P < 0.001).

**Figure 1.**
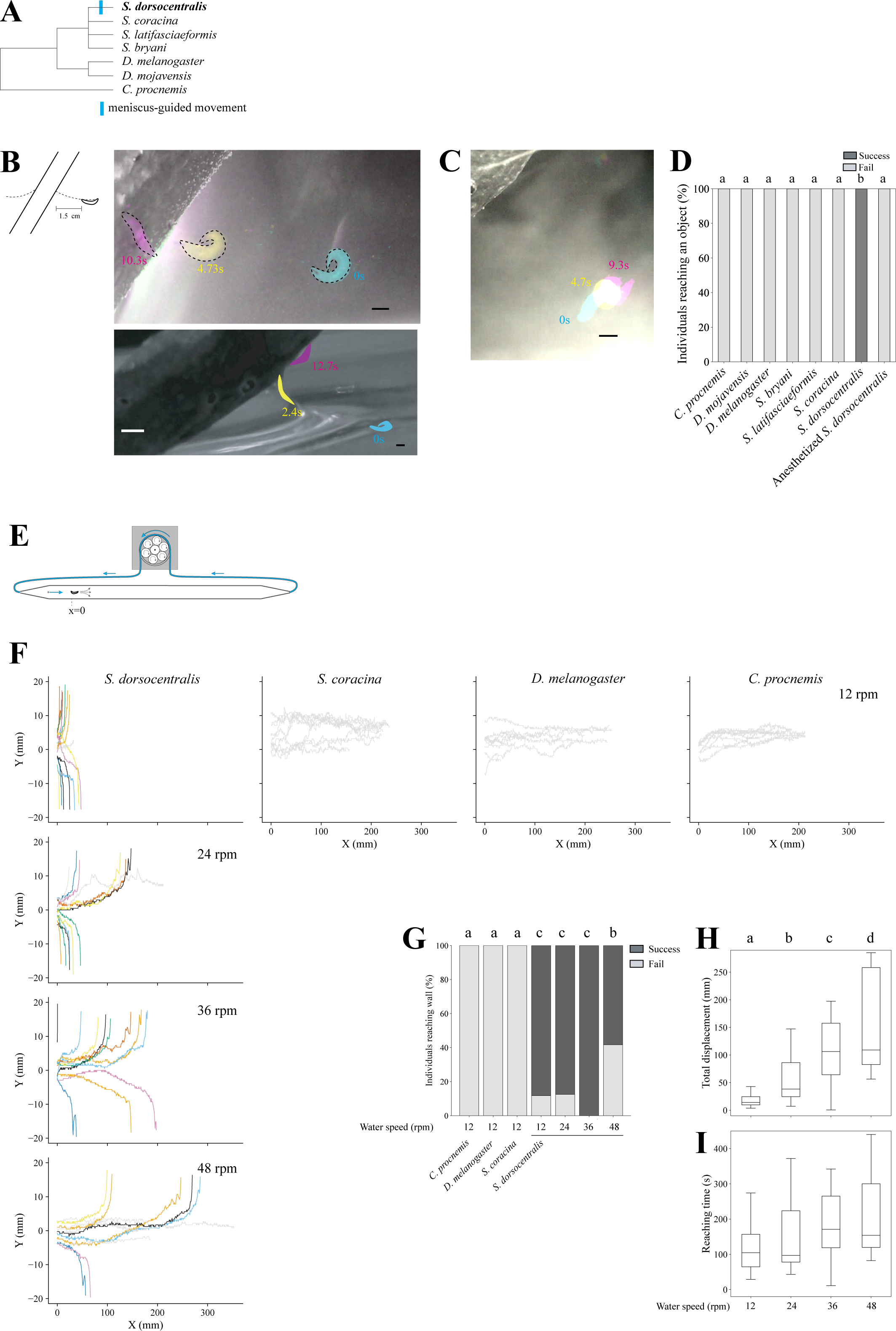
Larval *Scaptodrosophila dorsocentralis*, but not *Drosophila* or *Chymomyza*, moves along the water meniscus and reaches nearby objects under static and flowing conditions. (a) Phylogeny of Drosophilidae species used in this study. Branch lengths are not to scale. (b) Time-lapse images of *S. dorsocentralis* larvae placed on the water surface near a tree branch. The experimental setup is shown in the left panel; detailed dimensions are provided in Supplementary Fig. 1a. Dorsal (top) and lateral (bottom) views are shown. Pseudocolors indicate time progression. (c) Time-lapse images of *D. melanogaster* larvae under the same conditions. (d) Landing success rate (% of individuals reaching a nearby object). Successful individuals are shown in black and unsuccessful individuals in gray. P < 0.001, Fisher’s exact test with Holm–Bonferroni correction. Scale bars, 1 mm. N = 10-38 animals/treatment. (e) Schematic of the experimental setup. Detailed dimensions are provided in Supplementary Fig. 2. (f) Larval trajectories under different flow speeds. Each line represents an individual. Colored lines (non-gray) indicate individuals that successfully reached the wall within the 10 min recording period. Flow speed was controlled by adjusting the rotation rate of the peristaltic pump. According to the manufacturer’s specifications, a rotation speed of 48 rpm corresponds to approximately 29 mL min⁻¹. n= 10-17 animals/treatment (g) Proportion of individuals reaching the wall within 10 min. Successful individuals are shown in black and unsuccessful individuals in gray. Adjusted-P < 0.05 (a vs b), adjusted-P < 0.01 (a vs c), Fisher’s exact test with Holm–Bonferroni correction. (h) Total displacement along the x-axis under different flow speeds. Data were analyzed using a Kruskal–Wallis test followed by Mann–Whitney U tests with Holm–Bonferroni correction (adjusted P < 0.01 (a vs c) and P < 0.001 (a vs d), as indicated). (i) Reaching time to the arena wall under different flow speeds (Kruskal–Wallis test)

Together, these results demonstrate that *S. dorsocentralis* larvae perform directed movement along the water meniscus, a capability absent in closely related species. This behavior represents a previously unrecognized form of meniscus-guided movement in Diptera.

### Larval *S. dorsocentralis* moves along the water meniscus and reaches objects under flowing conditions

We next asked whether *S. dorsocentralis* can move along the water meniscus and reach objects under flowing conditions, mimicking semi-aquatic environments such as wind-driven or stream flow. To test this, we developed a behavioral arena (4 × 60 cm) in which larvae could move perpendicular to water flow generated by a peristaltic pump (Fig. 1e, Supplementary Fig. 1b). Chamber walls were lined with hydrophilic wooden material to create upward menisci. Larval movements on the water surface were recorded for 10 min. We tested four flow conditions (12, 24, 36, and 48 rpm), with 48 rpm corresponding to approximately 29 mL min⁻¹. At low flow (12 rpm), *S. dorsocentralis*, but not *S. coracina, D. melanogaster* or *C. procnemis*, moved perpendicular to the flow direction and rapidly reached the chamber walls (Fig. 1f, g; n = 10–17 animals per condition, Fisher’s exact test with Holm–Bonferroni correction, P < 0.01). Higher flow rates were tested only for *S. dorsocentralis*, as the other species failed to reach the wall even at 12 rpm. *S. dorsocentralis* maintained a high success rate: 88%–100% of individuals reached the wall at 12–36 rpm, and 58% succeeded even at the highest flow rate (48 rpm; Fig. 1f and g; n = 10–17 animals per condition; Fisher’s exact test with Holm–Bonferroni correction, P < 0.05). As flow speed increased, trajectories of *S. dorsocentralis* became increasingly displaced downstream, resulting in longer paths before reaching the wall (Fig. 1f,h; n = 10–17 animals per condition; Kruskal–Wallis test followed by Mann–Whitney U tests with Holm–Bonferroni correction). However, reaching time did not differ significantly across flow conditions (Fig. 1i). This indicates that the increased displacement at higher flow speeds reflects passive downstream drift rather than reduced efficiency of movement toward the wall.

Together, these results show that *S. dorsocentralis* retains the ability to perform meniscus-guided movement under flowing conditions, supporting the ecological relevance of this behavior and suggesting a capacity to function in water-associated environments.

### The species-specific behavioral sequence underlying S-shaped meniscus-guided movement

Because the S-shaped posture is specific to *S. dorsocentralis*, we next asked how this body configuration is generated and maintained at the air–water interface. To address this, we recorded larval behavioral sequences from a lateral view during the initiation and execution of meniscus-guided movement.

We observed that *S. dorsocentralis* larvae first elongate their posterior body while positioned at the air–water interface, with the posterior end anchored to the surface (Fig. 2a). The larvae then submerge their head into the water, forming a transient C-shaped posture, before rapidly elevating the head to generate an S-shaped body configuration (Fig. 2a,b). Once this posture is established, larvae move along the meniscus, initially at low speed and then gradually accelerating until they reach nearby objects or lose the S-shaped configuration (Fig. 2a,a′,b, Supplementary Movie 4).

**Figure 2.**
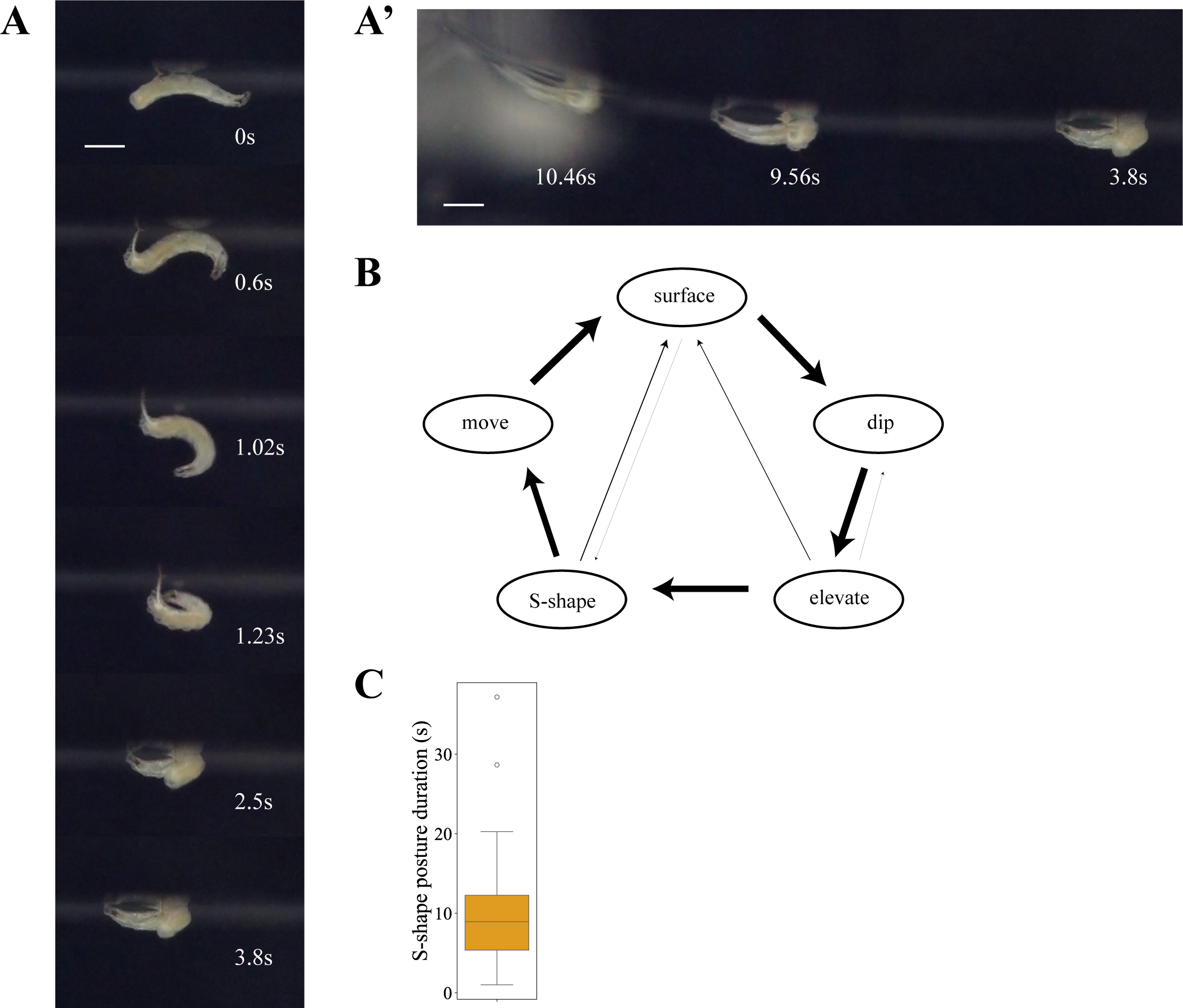
Behavioral sequence and maintenance of S-shaped body formation. (a) Time-lapse images of *S. dorsocentralis* larvae generating an S-shaped posture. With tail extended, the larva first dips its head (0.6 s), then elevates it (1.23 s), and subsequently forms an S-shape (2.5 s). (a′) The larva climbs the meniscus while maintaining the S-shaped posture. (b) State transition diagram of S-shaped body formation, comprising the states, flat, dip, elevate, S-shape, and move. Arrow thickness represents transition probability between states. The dominant transitions were surface → dip (0.97), dip → elevate (1.00), elevate → S-shape (0.89), and S-shape → move (0.85). A total of 161 transitions from 11 animals were analyzed. (c) Box plot showing the duration of the S-shaped posture following its formation. A total of 42 events from 9 animals were analyzed.

Transition probability analysis revealed that this behavioral sequence is highly stereotyped, with larvae progressing through a consistent series of states (Fig. 2b). In addition, the S-shaped posture was maintained for several seconds (median, 8.95 s), indicating that sustained posture maintenance is likely critical for effective meniscus climbing (Fig. 2c).

These behavioral sequences were not observed in other tested species, indicating that this strategy is specific to *S. dorsocentralis*.

### The Capillary forces generated by larvae are comparable to those in semi-aquatic insects

During formation of the S-shaped posture, larvae generate distinct meniscus deformations: upward menisci at the anterior tip and a middle region (approximately the A5 segment), and a downward meniscus at the hydrophobic posterior spiracle^25^ (Fig. 3a). This posture-dependent deformation of the water surface is reminiscent of previous reports where Hu and Bush (2005) established a physical framework for meniscus climbing based on torque balance, normal force balance, and tangential force balance in semi-aquatic insects^21,26^. We applied this framework to test whether the forces generated by *S. dorsocentralis* larvae are comparable to those reported in these systems.

**Figure 3.**
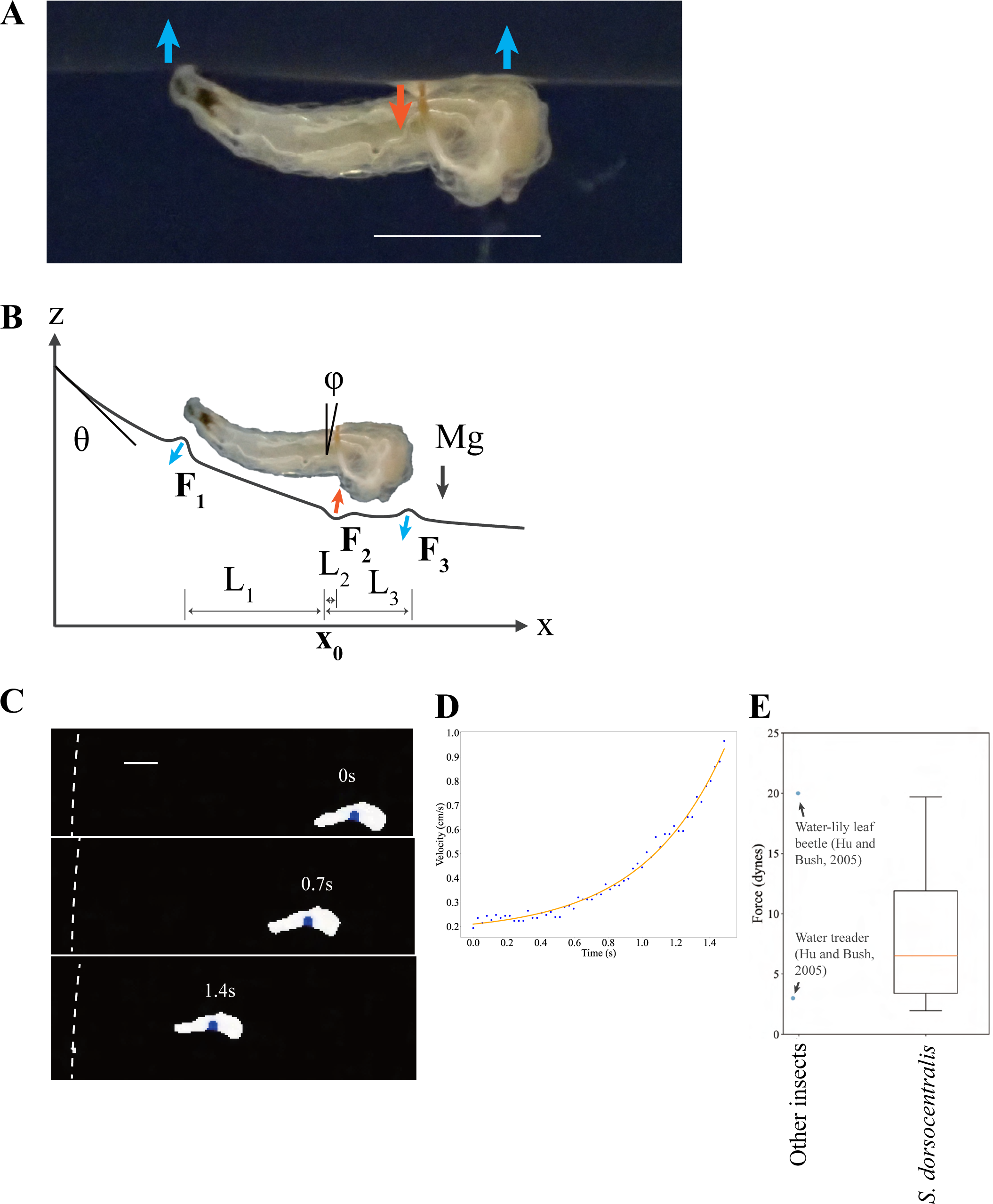
Estimation of capillary forces during meniscus climbing. (a) Schematic model of force generation. Surface tension forces are assumed to act at three points along the body: the anterior tip, a middle region (approximately the A5 segment), and the posterior tip. The anterior and middle points generate upward menisci, whereas the posterior point generates a downward meniscus, indicated by orange and cyan arrows, respectively. Parameters θ, L₁, L₂, L₃, φ, and Mg are defined in the figure. (b) Representative centroid trajectories of larvae in the Petri dish (blue). (c) Representative images of larvae approaching the Petri dish wall. The blue circles show centroids. Broken lines show the walls. (d) Representative velocity (cm s⁻¹) as a function of time (s). Blue dots indicate data points and the orange line shows the exponential fit. (e) Estimated forces (dynes). Forces reported for water lily beetles and water treaders are shown for comparison^63^ (n =22 animals). Scale bars, 1 mm.

We modeled capillary forces as acting at three points along the body, the anterior tip, a middle region, and the posterior tip, defined as 𝐹_1_, 𝐹_2_, and 𝐹_3_, respectively (Fig. 3b). To estimate these forces, we recorded larval movement along the meniscus and extracted centroid trajectories using LabGym^22^, following image preprocessing (binarization and filtering) in ImageJ (Fig. 3c, Supplementary Movie 5). From these trajectories, we calculated velocities and accelerations, and fitted acceleration as a function of displacement (Fig. 3d,e).

Additional parameters, including 𝐿_1_, 𝐿_2_, and 𝐿_3_, body mass, and contact angle, were measured independently and incorporated into the model. Solving the resulting equations yielded estimates of 𝐹_1_, the primary force driving movement toward the wall.

We found that the magnitude of 𝐹_1_is comparable to that reported for semi-aquatic insects. Specifically, the 10th–90th percentile of 𝐹_1_ranged from ∼3 to 20 dynes, overlapping with values reported for water lily leaf beetles (∼20 dynes) and water treaders (∼3 dynes)^21^. The variability observed in our measurements likely reflects differences in the efficiency of water-surface deformation among individuals.

These results indicate that *S. dorsocentralis* larvae generate capillary forces of similar magnitude to those of established semi-aquatic insects, supporting the idea that they exploit water-surface deformation to achieve meniscus-guided movement.

### PDMS-based phenocopy recapitulates larval meniscus-guided movement without external actuation

To test whether the S-shaped body configuration is sufficient to generate meniscus-guided movement, we constructed a physical mimic using a soft material. Polydimethylsiloxane (PDMS), commonly used in soft robotics, was molded and manually shaped to approximate the body form of *S. dorsocentralis* larvae. To reproduce the asymmetric body configuration, an insect pin was inserted to control the relative height of the anterior and posterior ends. Specifically, the posterior end was depressed to mimic the larval S-shaped posture, resulting in elevation of the anterior region (Fig. 4a).

**Figure 4.**
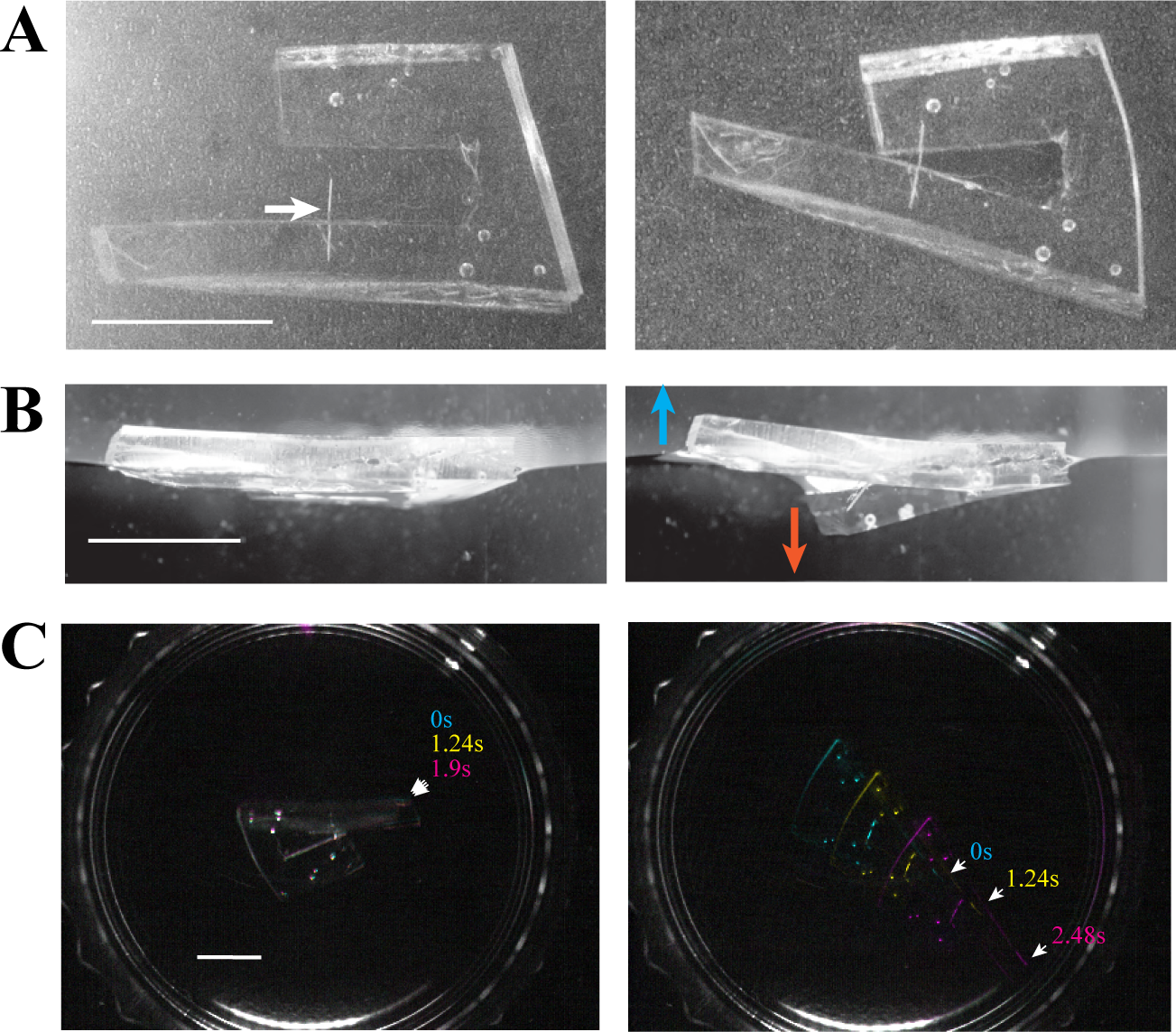
PDMS-based phenocopy of larval meniscus -guided movement without external actuation. (a) PDMS larval mimic. The left panel shows the unconnected structure, whereas the right panel shows the mimic after the anterior and posterior ends were connected using an insect pin. The white arrow indicates the insect pin. (b) Lateral views of PDMS constructs floating on the water surface. The horizontally connected control (left) generates minimal meniscus deformation, whereas the asymmetrically connected mimic (right), with the anterior end elevated and the posterior end depressed, generates upward and downward menisci, as indicated by arrows. (c) Time-lapse images of movement on the water surface. Pseudocolors indicate time progression. The left panel shows the control PDMS construct, and the right panel shows the asymmetrically connected mimic, both placed in a Petri dish containing 2 mL of water. Scale bars, 1 cm.

Under these conditions, the PDMS mimic generated clear meniscus deformations and moved along the water surface, ultimately reaching the wall of the Petri dish, recapitulating the movement of *S. dorsocentralis* larvae without any external actuation (Fig. 4). In contrast, control constructs with a horizontally aligned body, which produced minimal surface deformation, failed to move along the meniscus and did not reach the wall (Supplementary Movie 6 and 7).

These results demonstrate that the S-shaped body configuration is critical to generate capillary forces required for movement, supporting the idea that posture alone enables meniscus-guided movement in *S. dorsocentralis*.

### *Scaptodrosophila* larvae elongate their tails via a folding-based mechanism

Because an elongated posterior is required for the formation of the S-shaped posture, positioning the hydrophobic posterior end between the anterior and middle regions to generate asymmetric meniscus deformations (Fig. 3a), we hypothesized that ability to elongate the tail is a key factor contributing to meniscus-guided movement. We therefore analyzed tail elongation behavior in *Scaptodrosophila* larvae. To quantify tail elongation, we used a previously established dig-and-dive assay^23^, in which larvae were allowed to burrow into 0.5% agar while being recorded. Tail length at maximal extension was measured relative to total body length. We found that all tested *Scaptodrosophila* species display higher tail-to-body ratios than *D. melanogaster, D. mojavensis* and *C. procnemis* (n = 8–11 animals per species; Kruskal–Wallis test followed by Holm–Bonferroni correction, adjusted P < 0.001; Fig. 5a,b, Supplementary Movie 8). Notably, *D. mojavensis* exhibited higher tail-to-body ratios than *D. melanogaster* or *C. procnemis*, consistent with previous findings (adjusted P < 0.05; Fig. 5b) ^27^.

**Figure 5.**
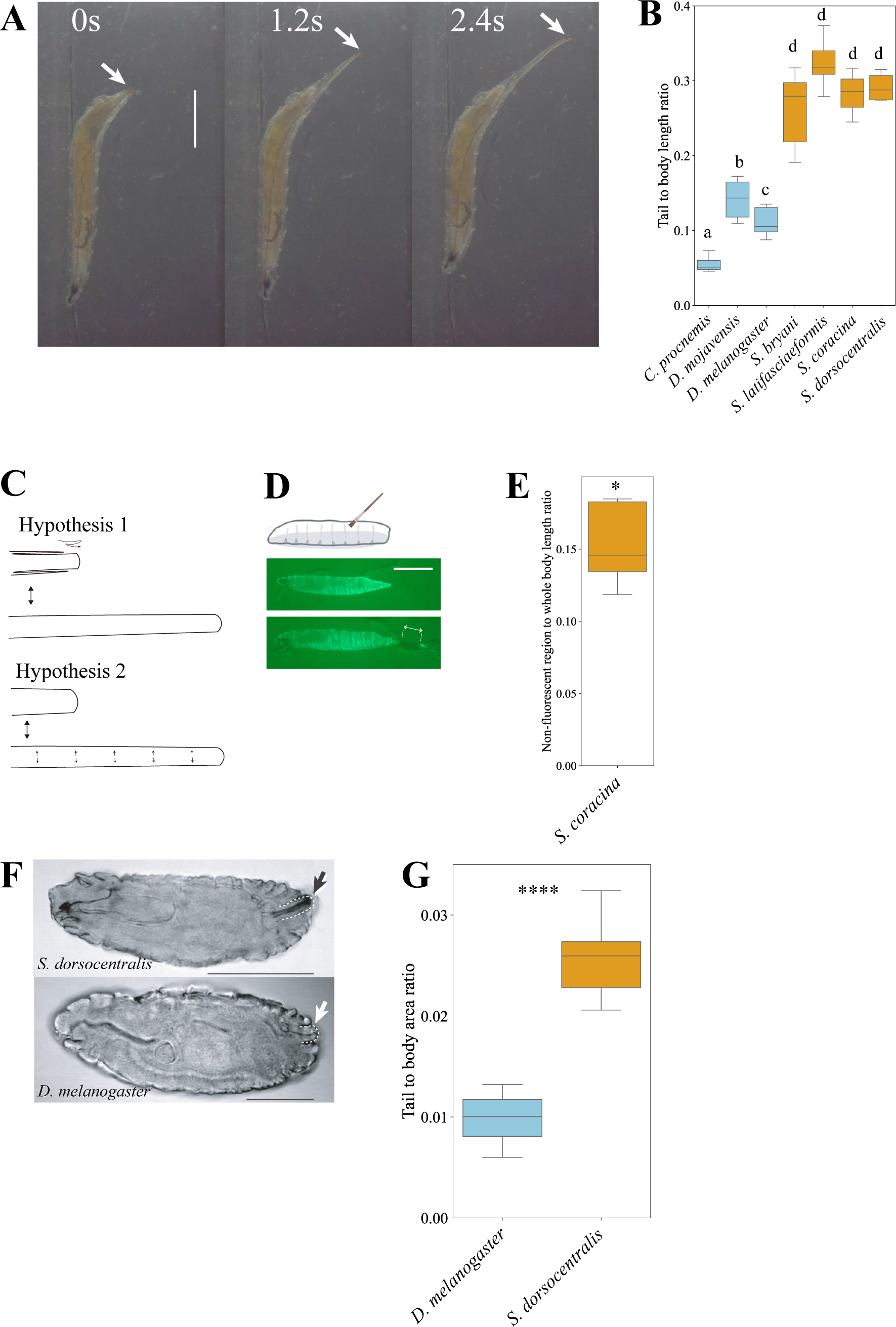
Tail elongation in *Scaptodrosophila* and its folding mechanism. (a) Time-lapse images of tail elongation in *S. coracina* larvae in 0.5% agar. (b) Comparison of tail-to-body length ratio across species (n = 8–11 animals/species). Kruskal–Wallis test followed by Mann–Whitney U tests with Holm–Bonferroni correction, adjusted P < 0.01 (a vs b, a vs c, a vs d, b vs d, or c vs d), adjusted P<0.05 (b vs c). (c) Two hypotheses for tail elongation. The upper panel illustrates the folding–unfolding model, whereas the lower panel illustrates the shrink–swell model. (d) Visualization of folded structures using fluorescent labeling. The top panel shows the schematic of dye application with a paintbrush. The middle panel shows the larva immediately after labeling, and the bottom panel shows the larva after tail elongation. (e) Ratio of the non-fluorescent axial region to total body length. Wilcoxon signed-rank test against a null hypothesis of median = 0 (n = 7 animals). (f) Stage 16 embryos of *S. dorsocentralis* (top) and *D. melanogaster* (bottom). White dashed lines indicate folded structures, which are more prominent in *S. dorsocentralis*. (g) Comparison of tail-to-body area ratio between *S. dorsocentralis* and *D. melanogaster* embryos (n = 15–17 animals/species). Mann–Whitney U test, ****P < 0.0001.

We next investigated the structural basis of tail elongation. We considered two alternative mechanisms: (1) a folding–unfolding mechanism, in which the tail contains a folded structure (filzkörper^28^) that expands upon extension, and (2) an elastic mechanism, in which the tissue stretches relatively uniformly (Fig. 5c). To distinguish between these models, we applied a fluorescent dye to larvae while the tail was in a contracted state and examined the distribution of the dye following elongation. Because *S. dorsocentralis* is small and difficult to assay, we used larger species *S. coracina* for this assay.

Upon tail extension, a distinct non-fluorescent region emerged, consistent with exposure of previously folded structures. This pattern supports the folding–unfolding model rather than uniform stretching (Fig. 5d,e; median ratio of non-fluorescent region length to total body length = 0.145, n = 7 animals, Wilcoxon signed-rank test against a null hypothesis of median = 0, P<0.05), although we cannot exclude a contribution from elastic deformation. These results indicate that tail elongation is achieved, at least in part, by deployment of a folded cuticular structure, particularly around the posterior spiracular region.

Then, we asked whether this folding structure is established during embryogenesis. We compared late-stage embryos between species and found that *S. dorsocentralis* embryos exhibit a more pronounced folded posterior structure than stage-matched *D. melanogaster* embryos (Fig. 5f,g; n = 15–17 animals per species; Mann–Whitney U test, P < 0.0001).

Together, these results indicate that the structural basis of tail elongation, which likely contributes to S-shape formation, arises during development in *Scaptodrosophila* and is mediated by a folding–unfolding mechanism.

### Posterior extension is likely driven by hydrostatic pressure generated by anterior–middle segment contraction

We next asked how the folded tail structure is unfolded during extension. We considered two alternative mechanisms: (1) contraction of the anterior-to-middle body segments generates hydrostatic pressure that drives unfolding of the posterior structure, and (2) tail elongation is driven by specialized transverse muscles in the posterior region (Fig. 6a^29–31^). Given that gain or loss of muscle structures can occur within Drosophilidae^32^, the second hypothesis represents a plausible lineage-specific mechanism.

**Figure 6.**
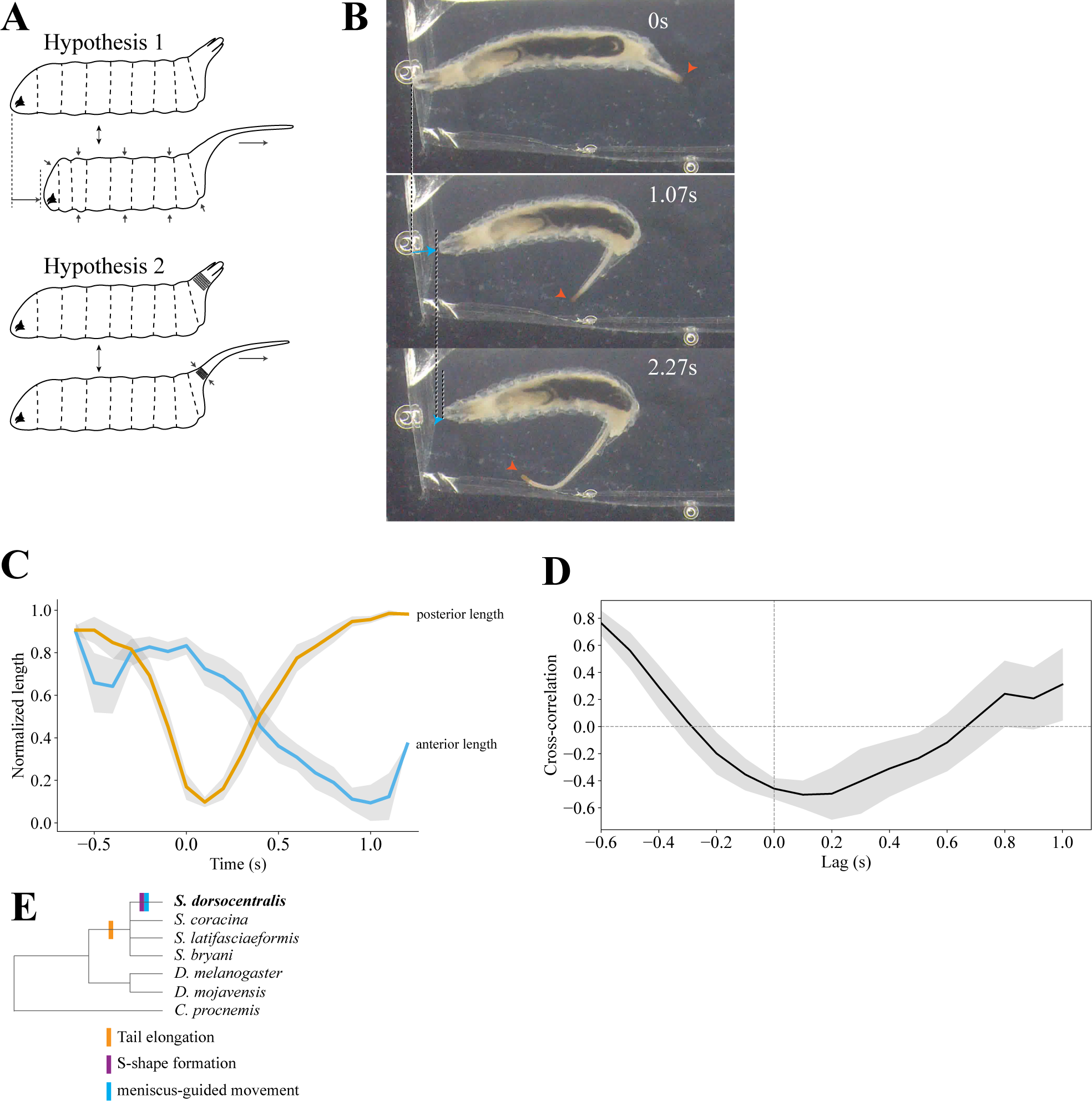
Tail elongation is driven by hydrostatic pressure generated by anterior–middle segment compression. (a) Two hypotheses for the power source of tail elongation. The upper panel illustrates local muscle contraction at the posterior end, whereas the lower panel illustrates compression of the anterior-to-middle segments generating hydrostatic pressure. (b) Representative time-lapse images of tail elongation in *S. coracina* larvae on agar. The gut was stained with carbon black for reference. Orange arrows indicate the posterior end, and blue arrows indicate the anterior end. (c) Posterior (orange) and anterior (cyan) lengths as a function of time (n = 14 animals). (d) Cross-correlation as a function of lag (s). Positive lag indicates that posterior elongation follows anterior contraction. (n = 14 animals). (e) Updated phylogeny showing the acquisition of tail elongation and S-shaped formation associated with meniscus-guided movement. These traits are indicated by rectangles along the lineage leading to *S. dorsocentralis*.

To test the first hypothesis, we analyzed segmental body dynamics by labeling the gut as a reference through feeding *S. coracina* larvae carbon black. Larvae were placed in 0.5% agar and their digging behavior was recorded. Anterior length was defined as the distance between the anterior tip and the A1/A2 segment boundary, and posterior length as the distance between the posterior tip and the A7/A8 segment boundary; lengths were normalized within individuals (Supplementary Fig. 3).

Analysis of anterior and posterior segment lengths revealed that contraction of the anterior-to-middle segments is temporally coupled with posterior extension (Fig. 6b,c; n = 14 animals, Supplementary Movie 9). Cross-correlation analysis revealed a broad negative correlation spanning lags from −0.3 to +0.6 s, indicating sustained anti-phase coupling between anterior and posterior body regions. (Fig. 6d; n = 14 animals). This pattern is consistent with hydrostatic pressure-mediated coupling, where contraction in one region leads to expansion in another.

To test the second hypothesis, we examined whether posterior-specific musculature could account for tail elongation. Phalloidin staining revealed that the major transverse muscle groups, whose contractions are expected to drive posterior unfolding, are broadly conserved between *D. melanogaster* and *S. coracina* (Supplementary Fig. 2; n = 5 animals per species). This suggests that tail elongation is unlikely to be driven by lineage-specific muscle architecture, although finer-scale analyses will be required to fully exclude this possibility.

Together, these results support a model in which contraction of the anterior-to-middle body segments increases hydrostatic pressure, leading to deployment of the folded posterior structure. This mechanism likely represents a physical basis of tail elongation that is co-opted in *S. dorsocentralis* for S-shaped posture formation (Fig. 6e).

## DISCUSSION

In this study, we show that larvae of *Scaptodrosophila dorsocentralis* can move along the water meniscus and reach nearby objects. We further demonstrate that this movement depends on the formation of an S-shaped body posture, which can be recapitulated using a PDMS-based physical mimic, and that tail elongation is achieved through a folding-based morphological mechanism coupled with anterior–middle segment compression. Together, these findings reveal a previously unrecognized form of meniscus-guided movement in Diptera and provide a mechanistic framework linking body posture, morphology, and physical forces at the air–water interface.

Although some *Scaptodrosophila* species have been noted as having a particular affinity to moist environments ^33,34^, the ecology of *S. dorsocentralis* remains largely unknown. The species was originally collected in Okinawa, a subtropical environment where water is abundant^17^, but it is unclear whether larvae naturally encounter aquatic or semi-aquatic conditions. Nevertheless, meniscus climbing has been reported across diverse insects, including semi-aquatic species, such as water lily leaf beetles and water treaders^13,14,21^. In our study, the ability of *S. dorsocentralis* larvae to reach nearby objects via the water meniscus was maintained even under moderate flow conditions, supporting the ecological relevance of this behavior. These observations raise the possibility that *S. dorsocentralis* occupies, or is transitioning toward, a semi-aquatic niche. However, direct ecological validation through field observations will be required, although identifying larvae in natural environments may be challenging due to their small size and cryptic habitats.

Transitions between terrestrial and freshwater environments have occurred repeatedly in insects and are a major driver of biodiversity. Although freshwater habitats harbor a disproportionate fraction of global diversity^1^, the mechanisms by which terrestrial insects adapt to these environments remain poorly understood^9,10^. Diptera, in particular, includes a large number of lineages that exploit aquatic habitats at some stage of their life cycle^1,3,15,35^. Indeed, larvae of a few drosophilid species (for example, the *D. simulivora* group) have been reported to be fully aquatic, and closely related lineages such as Ephydridae (shore flies) have extensively adapted to aquatic or semi-aquatic environments ^24,36–38^. The discovery of meniscus-guided movement in *S. dorsocentralis* therefore suggests that some drosophilids possess latent capacities for exploiting water-surface environments and may represent an early stage in the transition from terrestrial to semi-aquatic niches. As such, *S. dorsocentralis* provides a promising model system for investigating the mechanisms underlying niche transitions.

A key morphological feature associated with this behavior is tail elongation. We show that *Scaptodrosophila* larvae extend their posterior body via a folding–unfolding mechanism driven, at least in part, by hydrostatic pressure generated through anterior–middle segment compression. Similar elongate posterior structures are found in other semi-aquatic dipteran larvae, such as Ephydridae and rat-tailed maggots (Eristalini, Syrphidae), where they function as respiratory siphons^38,39^. Although the functional roles differ, the repeated evolution of elongated posterior structures in these lineages suggests a potential link between posterior extensibility and adaptation to water-associated environments. It is therefore possible that tail elongation in *Scaptodrosophila* represents an early or intermediate state in the evolution of semi-aquatic adaptations within Cyclorrhapha, and potentially more broadly across Diptera^40^.

Our results also highlight the importance of physical interactions between the body and the environment. By combining behavioral analysis, physical modeling, and biomimetic reconstruction, we show that an S-shaped posture, in which the hydrophobic posterior spiracle generates a downward meniscus and is positioned between upward menisci formed at the anterior and middle regions, enables larvae to produce asymmetric surface deformations and capillary forces sufficient for movement. Notably, the magnitude of these forces is comparable to those reported for established semi-aquatic insects, despite the largely terrestrial ecology of drosophilids. This suggests that relatively simple changes in body configuration and mechanical coupling to the environment can give rise to novel movement strategies through the exploitation of physical forces, without requiring extensive morphological specialization.

Several important questions remain. First, the genetic and developmental mechanisms underlying tail folding and extensibility are currently unknown. Although our results indicate that the folded structure is already established during embryogenesis in *Scaptodrosophila*, the gene regulatory networks that govern its formation and evolutionary modification remain to be elucidated^28,41^. Notably, this behavior is not observed in other tested *Scaptodrosophila* species, which exhibit tail elongation but lack meniscus-guided movement, suggesting that the neural circuits required for coordinating this behavior are absent or functionally divergent in these species (Fig. 6 e). In particular, it will be important to understand how larvae sense the air–water interface and coordinate motor outputs to maintain a stable configuration over several seconds. One possibility is that neuroendocrine or neuromodulatory pathways involved in osmoregulation contribute to this process. For example, Leucokinin signaling, which regulates water balance in insects, could be activated upon contact with the water surface^42–46^. Alternatively, sensory pathways detecting humidity or moisture gradients may be involved. In insects, Ionotropic receptors ^47–49^, and Odorant receptors ^50^ are known to mediate hygrosensation and could contribute to detecting the air–water interface. Maintaining the S-shaped posture is also likely to require precise proprioceptive feedback. In *Drosophila* larvae, genes such as *tmc*, *pzl*, and *ninaE* have been implicated in proprioceptive control of body posture ^51–55^, suggesting that similar mechanisms may contribute to posture stabilization in *S. dorsocentralis*. These mechanisms may represent species-specific innovations or may arise through co-option of ancestral sensory and motor circuits. Addressing these questions will require the development of functional genetic tools in *Scaptodrosophila*.

Finally, our study establishes *S. dorsocentralis* as a promising system for investigating the emergence of novel behaviors at the interface of physics and evolution^56–66^. As a close relative of *D. melanogaster*, *Scaptodrosophila* offers the potential to integrate genetic, neurobiological, and physical approaches in a tractable framework. More broadly, our findings suggest that interactions with fluid interfaces may provide a rich and underexplored axis for the evolution of behavior and morphology.

## ACKNOWLEDGEMENTS

We thank the Kyoto Drosophila Stock Center, the National Drosophila Stock Center (Cornell University), and the Kyorin-Fly Drosophila Species Stock Center for providing fly strains. We are grateful to Takeshi Awasaki for technical advice on culturing *Scaptodrosophila*, to Hiroshi Kohsaka for advice on behavioral assay setup, and to Bernard Kim for assistance with phylogenetic analysis. We thank Maarten Zwart, Katrin Vogt, Armin Bahl, Yuma Tsukasa, and Tadao Usui for helpful discussions. We also thank Masaya Takahashi, Takahisa Date, Takaki Seki, and Tomohiro Kubo for assistance with fly maintenance, and members of the Nose laboratory for fly food preparation, as well as helpful discussions and feedback. This work was supported by the Japan Society for the Promotion of Science (grant no. 25K09722 to T.M.) and the Toyota Foundation (grant no. D24-HS-0144 to T.M.).

## FIGURE SUPPLEMENT CAPTIONS

**Supplementary figure 1: Schematic representations of behavioral assays.** (a) Schematic of the larval landing assay under static water conditions. A tree branch (*Zelkova serrata*; 2 cm diameter, 7 cm length) collected on campus was submerged in a cubic glass chamber filled with tap water. A larva was placed on the water surface at a distance of ∼1.5 cm from the branch. (b) Schematic of the larval landing assay under flowing water conditions. A peristaltic pump was used to generate variable flow rates. A larva was placed on the water surface and recorded for up to 10 min or until it reached the chamber wall (1 fps). The chamber walls were lined with thin hydrophilic wooden boards. Dorsal view. (b′) Lateral view of the flowing water assay. Slight slopes were introduced at the inlet and outlet to maintain continuous flow. (c) Schematic of the assay used for estimating capillary forces. Larval movements were recorded at 37 fps.

**Supplementary Figure 2 No gross differences in transverse muscle organization in the posterior region between *S. coracina* and *D. melanogaster* larvae**

(a) Phalloidin conjugated to Alexa Fluor 488 was used to stain muscles in third instar *D. melanogaster* larvae. Images were acquired using a 4× air objective. Arrowheads indicate three transverse muscles in the posterior region. (b–b′) Phalloidin–Alexa Fluor 488 staining of third instar *S. coracina* larvae. Images were acquired using a 4× air objective (b) or a 20× objective (b′). Arrowheads indicate three transverse muscles in the posterior region. Scale bars, 100 µm.

**Supplementary Figure 3 Definition of posterior and anterior lengths**

Representative image of *S. coracina* larvae undergoing tail elongation. The magenta polygon indicates the posterior length, and the green polygon indicates the anterior length.

## SUPPLEMENTARY MATERIAL AND METHODS

### Body wall muscles staining

The protocol was adapted from Matsunaga et al. (2016)^67^. Briefly, third instar larvae were dissected in PBS to prepare larval fillets on Sylgard-coated plates (Sylgard 184, Dow) using fine forceps, microscissors (LMB-54-1, Nazme) and insect pins. Samples were fixed in 4% formaldehyde for 15 min, followed by rinsing in PBT (PBS with 0.1% Triton X-100). Samples were then blocked in 10% normal goat serum in PBT. For muscle staining, samples were incubated overnight at 4 °C with Phalloidin–FITC (diluted in blocking solution). After rinsing in PBT, larval musculature was imaged using a confocal microscope (FV3000, Olympus) with a 488 nm laser, using either a 4× air objective (UPLANSApo, Olympus) or a 20× objective (XLUMPlanFL N, Olympus).

## SUPPLEMENTARY INFORMATION

**Supplementary Movie 1:** *S. dorsocentralis* larvae move along the water meniscus, reach nearby objects, and land on them. Dorsal view.

**Supplementary Movie 2:** *S. dorsocentralis* larvae move along the water meniscus, reach nearby objects, and land on them. Positions are indicated by white crosses. Playback speed, 0.5×. Lateral view.

**Supplementary Movie 3:** *D. melanogaster* larvae placed on the water surface exhibit uncoordinated movements without directional displacement.

**Supplementary Movie 4:** Formation and maintenance of the S-shaped posture in *S. dorsocentralis* larvae at the air–water interface. Lateral view.

**Supplementary Movie 5:** Movement of *S. dorsocentralis* larvae in a Petri dish filled with water. Centroids are indicated by blue circles.

**Supplementary Movie 6:** Control PDMS construct exhibiting minimal meniscus deformation in a Petri dish filled with water. The anterior end is indicated by white crosses. Playback speed, 0.5×.

**Supplementary Movie 7:** Experimental PDMS mimic exhibiting asymmetric meniscus deformation in a Petri dish filled with water. The anterior end is indicated by white crosses. Playback speed, 0.5×.

**Supplementary Movie 8:** Tail elongation of *S. coracina* larvae in 0.5% agar.

**Supplementary Movie 9:** Tail elongation of *S. coracina* larvae in 0.5% agar is accompanied by anterior body compression. The gut is visualized using carbon black–supplemented food.

## REFERENCES

1. Dijkstra, K.-D. B., Monaghan, M. T. & Pauls, S. U. Freshwater biodiversity and aquatic insect diversification. Annu. Rev. Entomol. 59, 143–163 (2014).

2. Anderson, N. H. & Wallace. Habitat, life history, and behavioral adaptations of aquatic insects. (2010).

3. Adler, P. H. & Courtney, G. W. Ecological and societal services of aquatic Diptera. Insects 10, 70 (2019).

4. Matsunaga, T. et al. Odorant receptors mediating avoidance of toxic mustard oils in Drosophila melanogaster are expanded in herbivorous relatives. Mol. Biol. Evol. (2025) doi:10.1093/molbev/msaf164.

5. Matsunaga, T. et al. Evolution of Olfactory Receptors Tuned to Mustard Oils in Herbivorous Drosophilidae. Mol. Biol. Evol. 39, msab362 (2021).

6. Bush, J. W. M., Hu, D. L. & Prakash, M. The integument of water-walking arthropods: Form and function. in Advances in Insect Physiology (eds. Casas, J. & Simpson, S. J.) vol. 34 117–192 (Elsevier, 2007).

7. Watson, D. A. et al. Water striders are impervious to raindrop collision forces and submerged by collapsing craters. Proc. Natl. Acad. Sci. U. S. A. 121, e2315667121 (2024).

8. Dickerson, A. K., Shankles, P. G., Madhavan, N. M. & Hu, D. L. Mosquitoes survive raindrop collisions by virtue of their low mass. Proc. Natl. Acad. Sci. U. S. A. 109, 9822–9827 (2012).

9. Zhang, Q.-L. et al. Comparative transcriptomic analysis of fireflies (Coleoptera: Lampyridae) to explore the molecular adaptations to fresh water. Mol. Ecol. 29, 2676–2691 (2020).

10. Almudi, I. et al. Genomic adaptations to aquatic and aerial life in mayflies and the origin of insect wings. Nat. Commun. 11, 2631 (2020).

11. Misof, B. et al. Phylogenomics resolves the timing and pattern of insect evolution. Science 346, 763–767 (2014).

12. Bush, J. W. M. & Hu, D. L. WALKING ON WATER: Biolocomotion at the interface. Annu. Rev. Fluid Mech. 38, 339–369 (2006).

13. Ortega-Jiménez, V. M., Arriaga-Ramirez, S. & Dudley, R. Meniscus ascent by thrips (Thysanoptera). Biol. Lett. 12, (2016).

14. Miyamoto, S. On a special mode of locomotion utilizing surface tension at the water-edge in some semiaquatic insects. Kontyû 23, 45–52 (1955).

15. Sundermann, A., Lohse, S., Beck, L. A. & Haase, P. Key to the larval stages of aquatic true flies (Diptera), based on the operational taxa list for running waters in Germany. Ann. Limnol. 43, 61–74 (2007).

16. Jennings, B. H. Drosophila – a versatile model in biology & medicine. Mater. Today (Kidlington) 14, 190–195 (2011).

17. Okada, T. DROSOPHILIDAE OF THE OKINAWA ISLANDS. Kontyu (1965).

18. Wheeler, M. R. The Drosophilidae of the Nearctic Region exclusive of the genus Drosophila. The University of Texas Publication 5204, (1952).

19. Kim, B. Y. et al. Single-fly genome assemblies fill major phylogenomic gaps across the Drosophilidae Tree of Life. PLoS Biol. 22, e3002697 (2024).

20. Kim, B. Y. et al. Highly contiguous assemblies of 101 drosophilid genomes. Elife 10, (2021).

21. Hu, D. L. & Bush, J. W. M. Meniscus-climbing insects. Nature 437, 733–736 (2005).

22. Hu, Y. et al. LabGym: Quantification of user-defined animal behaviors using learning-based holistic assessment. *Cell Rep*. Methods 3, 100415 (2023).

23. Kim, D., Alvarez, M., Lechuga, L. M. & Louis, M. Species-specific modulation of food-search behavior by respiration and chemosensation in Drosophila larvae. Elife 6, (2017).

24. Bächli, G. & Tsacas, L. Two new species of the Drosophila simulivora group (Diptera, Drosophilidae). Preprint at http://www.drosophila.jp/jdd/class/070101/07010104.pdf.

25. Manning, G. Development of the Drosophila tracheal system. The Development of Drosophila melanogaster 1, 609–685 (1993).

26. Chan, D. Y. C., Henry, J. D., jr & White, L. R. The interaction of colloidal particles collected at fluid interfaces. J. Colloid Interface Sci. 79, 410–418 (1981).

27. Kalay, G. et al. Evolution of larval segment position across 12 Drosophila species. Evolution 74, 1409–1422 (2020).

28. Hu, N. & Castelli-Gair, J. Study of the posterior spiracles of Drosophila as a model to understand the genetic and cellular mechanisms controlling morphogenesis. Dev. Biol. 214, 197–210 (1999).

29. Kohsaka, H., Okusawa, S., Itakura, Y., Fushiki, A. & Nose, A. Development of larval motor circuits in Drosophila. Dev. Growth Differ. 54, 408–419 (2012).

30. Liu, Y., Hasegawa, E., Nose, A., Zwart, M. F. & Kohsaka, H. Synchronous multi-segmental activity between metachronal waves controls locomotion speed in Drosophila larvae. Elife 12, e83328 (2023).

31. Zwart, M. F. et al. Selective inhibition mediates the sequential recruitment of motor pools. Neuron 91, 615–628 (2016).

32. Gailey, D. A. et al. The muscle of lawrence in Drosophila: a case of repeated evolutionary loss. Proc. Natl. Acad. Sci. U. S. A. 94, 4543–4547 (1997).

33. Parsons, P. A. Adaptive radiation in the subgenus Scaptodrosophila of australian Drosophila. Nature 258, 602 (1975).

34. Bock, I. R. & Parsons, P. A. The subgenus *Scaptodrosophila* (Diptera: Drosophilidae). Syst. Entomol. 3, 91–102 (1978).

35. Suzuki, C. et al. Hydrophobic-hydrophilic crown-like structure enables aquatic insects to reside effectively beneath the water surface. *Commun*. Biol. 4, 708 (2021).

36. Disney, R. H. L. Drosophila gibbinsi larvae also eat simulium. Trans. R. Soc. Trop. Med. Hyg. 69, 365–366 (1975).

37. Werner, D. & Pont, A. C. Dipteran predators of Simuliid blackflies: a worldwide review. Med. Vet. Entomol. 17, 115–132 (2003).

38. Cash-Clark, C. E. & Bradley, T. J. External morphology of the larvae of Ephydra (Hydropyrus) hians (Diptera: Ephydridae). J. Morphol. 219, 309–318 (1994).

39. Pfiester, M. & Kaufman, P. E. Drone fly, rat-tailed maggot Eristalis tenax (Linnaeus) (Insecta: Diptera: Syrphidae): EENY 445/IN809, 3/2009. EDIS 2009, (2009).

40. Wiegmann, B. M. et al. Episodic radiations in the fly tree of life. Proc. Natl. Acad. Sci. U. S. A. 108, 5690–5695 (2011).

41. Molina-Gil, S., Sotillos, S., Espinosa-Vázquez, J. M., Almudi, I. & Hombría, J. C.-G. Interlocking of co-opted developmental gene networks in Drosophila and the evolution of pre-adaptive novelty. Nat. Commun. 14, 5730 (2023).

42. Nässel, D. R. & Wu, S.-F. Leucokinins: Multifunctional neuropeptides and hormones in insects and other invertebrates. Int. J. Mol. Sci. 22, 1531 (2021).

43. Yu, M.-J. & Beyenbach, K. W. Effects of leucokinin-VIII on Aedes Malpighian tubule segments lacking stellate cells. J. Exp. Biol. 207, 519–526 (2004).

44. Al-Anzi, B. et al. The leucokinin pathway and its neurons regulate meal size in Drosophila. Curr. Biol. 20, 969–978 (2010).

45. Puri, S., Sang, J., Pandey, P. & Lee, Y. Insulin and leucokinin pathways coordinate adaptive salt appetite in Drosophila. Proc. Natl. Acad. Sci. U. S. A. 123, e2530544123 (2026).

46. Li, K. et al. Belly roll, a GPI-anchored Ly6 protein, regulates Drosophila melanogaster escape behaviors by modulating the excitability of nociceptive peptidergic interneurons. Elife 12, e83856 (2023).

47. Knecht, Z. A. et al. Ionotropic Receptor-dependent moist and dry cells control hygrosensation in Drosophila. Elife 6, e26654 (2017).

48. Enjin, A. et al. Humidity sensing in Drosophila. Curr. Biol. 26, 1352–1358 (2016).

49. Chu, L.-A., Tai, C.-Y. & Chiang, A.-S. Thirst-driven hygrosensory suppression promotes water seeking in Drosophila. Proc. Natl. Acad. Sci. U. S. A. 121, e2404454121 (2024).

50. Li, S., Li, B., Gao, L., Wang, J. & Yan, Z. Humidity response in Drosophila olfactory sensory neurons requires the mechanosensitive channel TMEM63. Nat. Commun. 13, 3814 (2022).

51. Guo, Y. et al. Transmembrane channel-like (tmc) gene regulates Drosophila larval locomotion. Proc. Natl. Acad. Sci. U. S. A. 113, 7243–7248 (2016).

52. He, L. et al. Direction selectivity in Drosophila proprioceptors requires the mechanosensory channel Tmc. Curr. Biol. 29, 945–956.e3 (2019).

53. Vaadia, R. D. et al. Characterization of proprioceptive system dynamics in behaving Drosophila larvae using high-speed volumetric microscopy. Curr. Biol. 29, 935–944.e4 (2019).

54. Zanini, D. et al. Proprioceptive opsin functions in Drosophila larval locomotion. Neuron 98, 67–74.e4 (2018).

55. Heckscher, E. S. et al. Even-skipped(+) interneurons are core components of a sensorimotor circuit that maintains left-right symmetric muscle contraction amplitude. Neuron 88, 314–329 (2015).

56. Tanaka, R. et al. Cross-species implementation of an innate courtship behavior by manipulation of the sex-determinant gene. Science 389, 747–752 (2025).

57. Auer, T. O. et al. Olfactory receptor and circuit evolution promote host specialization. Nature 579, 402–408 (2020).

58. Ishikawa, A. et al. A key metabolic gene for recurrent freshwater colonization and radiation in fishes. Science 364, 886–889 (2019).

59. Metz, H. C., Bedford, N. L., Pan, Y. L. & Hoekstra, H. E. Evolution and genetics of precocious burrowing behavior in Peromyscus mice. Curr. Biol. 27, 3837–3845.e3 (2017).

60. Seeholzer, L. F., Seppo, M., Stern, D. L. & Ruta, V. Evolution of a central neural circuit underlies Drosophila mate preferences. Nature 559, 564–569 (2018).

61. Young, L. J., Nilsen, R., Waymire, K. G., MacGregor, G. R. & Insel, T. R. Increased affiliative response to vasopressin in mice expressing the V1a receptor from a monogamous vole. Nature 400, 766–768 (1999).

62. Bendesky, A. et al. The genetic basis of parental care evolution in monogamous mice. Nature 544, 434–439 (2017).

63. Elipot, Y., Hinaux, H., Callebert, J. & Rétaux, S. Evolutionary shift from fighting to foraging in blind cavefish through changes in the serotonin network. Curr. Biol. 23, 1–10 (2013).

64. Prieto-Godino, L. L. et al. Evolution of acid-sensing olfactory circuits in drosophilids. Neuron 93, 661–676.e6 (2017).

65. Sakurai, A. & Katz, P. S. Artificial synaptic rewiring demonstrates that distinct neural circuit configurations underlie homologous behaviors. Curr. Biol. 27, 1721–1734.e3 (2017).

66. Bumbarger, D. J., Riebesell, M., Rödelsperger, C. & Sommer, R. J. System-wide rewiring underlies behavioral differences in predatory and bacterial-feeding nematodes. Cell 152, 109–119 (2013).

67. Matsunaga, T., Kohsaka, H. & Nose, A. Gap junction-mediated signaling from motor neurons regulates motor generation in the central circuits of larval Drosophila. J. Neurosci. 37, 2045–2060 (2017).

