## Supplementary figures and images for "Emergence of meniscus-guided movement in drosophilid larvae through posture-dependent capillary forces"

### Supplementary Figs

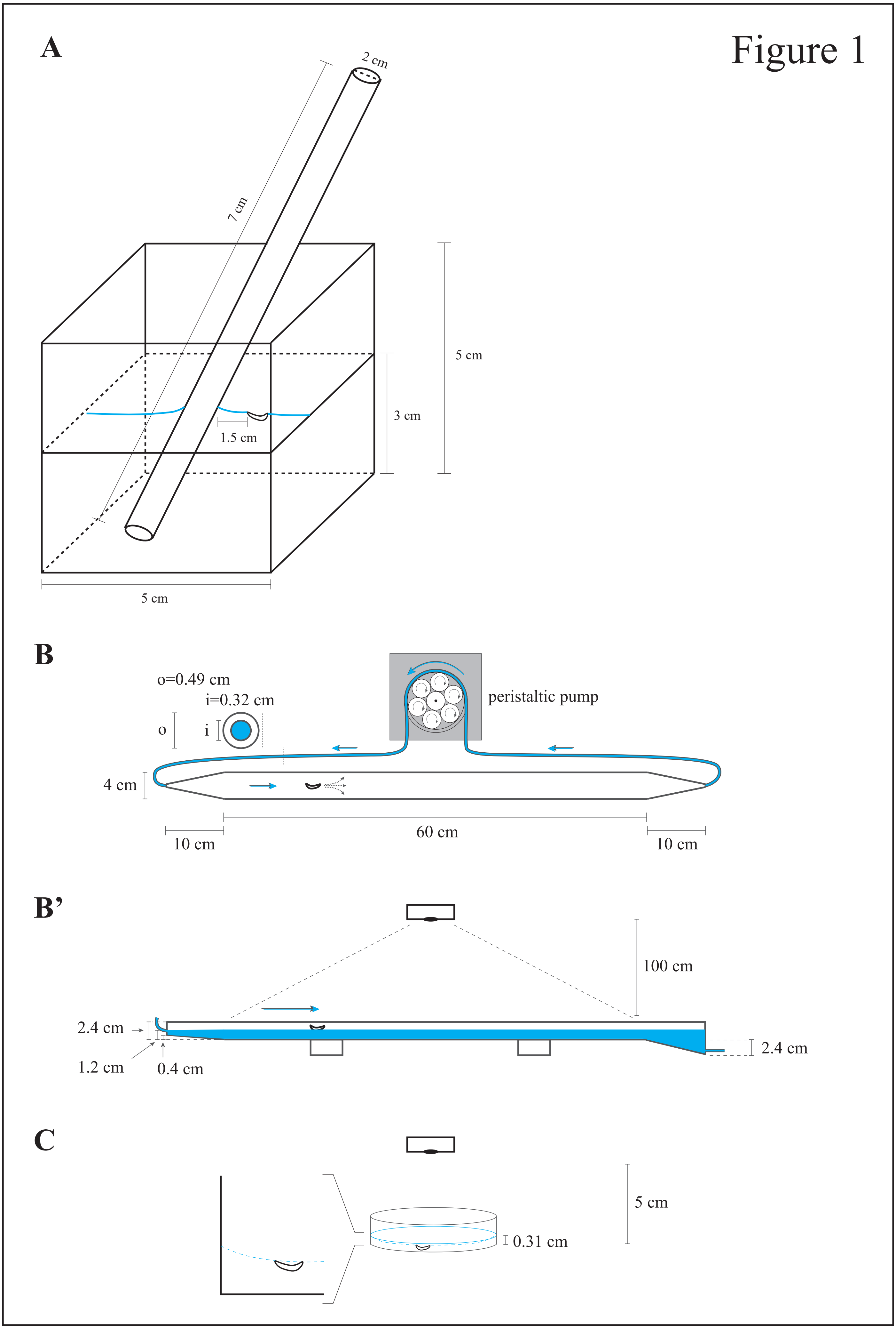

Figure 2

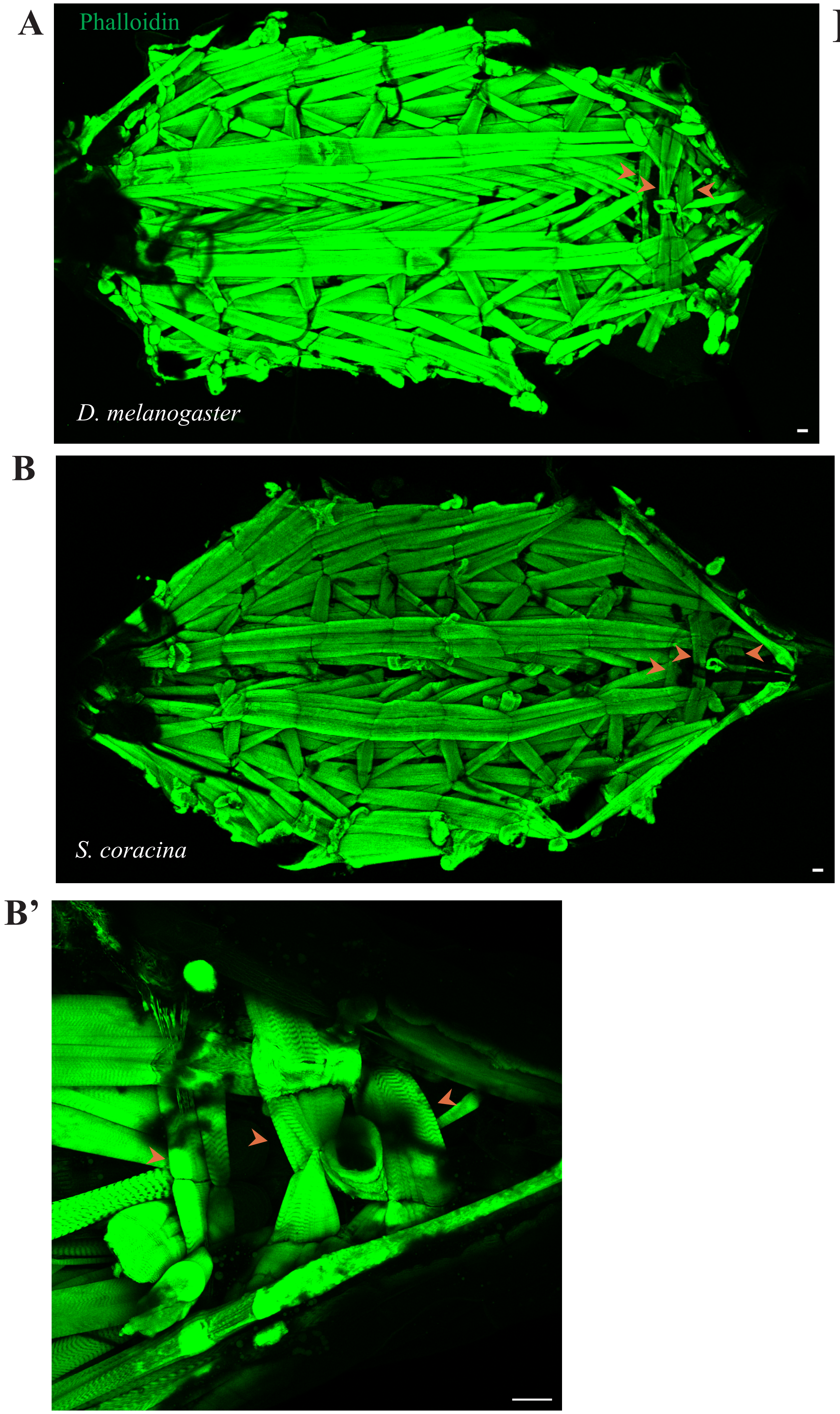

Figure 3

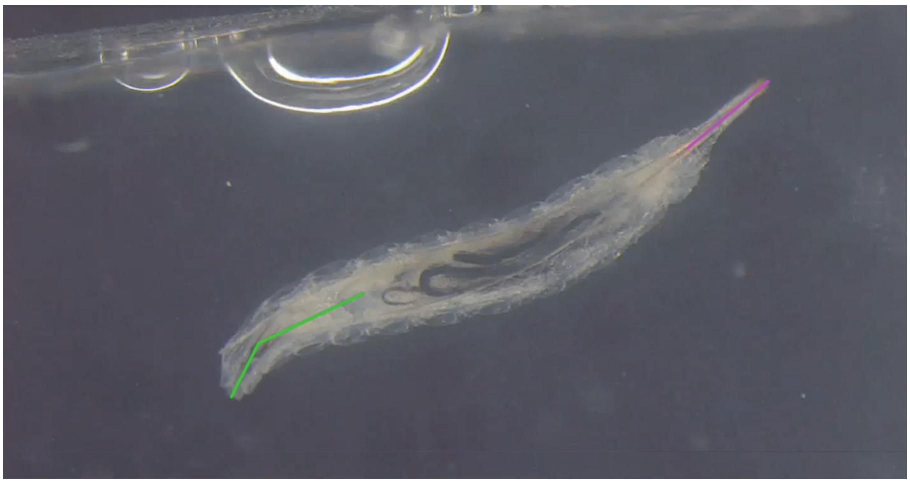
